# Quantum-like dynamics in the human brain

**DOI:** 10.1101/2025.10.02.680057

**Authors:** Gustavo Deco, Yonatan Sanz Perl, Natasha Greenstein, Shamil Chandaria, Greg Scholes, Morten L. Kringelbach

## Abstract

Emerging new research indicates evidence of quantum-like (QL) probability laws, including interference effects, in non-quantum physical systems using coupled oscillators. This can produce QL states which can compute in a QL fashion. Given the success of using coupled oscillators for human whole-brain modelling, we investigate the possibility of QL dynamics in the human brain. Here, we investigate how the special topology of human brain anatomy together with QL bits can promote the rich dynamic repertoire necessary for human advanced cognition. We systematically changed the level of QL processing in a whole-brain model. We found the QL regime provided the best whole-brain model fit to large-scale human empirical neuroimaging data. Extraordinarily, at this optimum point we found significantly lower energy consumption than for the non-QL networks. Mechanistically, this implies that the significantly larger whole-brain spectral gap for QL networks offers a backbone to the functional metastability needed to provide the necessary dynamical regime for efficient computation. The underlying QL spectral gaps amplify through interference the metastability and richness of repertoire of the human brain. Overall, we found that the special topology of the human brain promotes QL information processing.

## Introduction

The human brain shows remarkable computational powers on a very limited energy budget [1, 2]. Somehow this must be achieved through distributed, efficient time and energy computation, requiring long-range interactions as suggested by the unique human brain architecture with a preponderance of rare long-range exceptions to the exponential distance rule [3–6]. These underlying mechanisms must also deal with the relative slowness of information processing through neurons. Overall, this poses a major unsolved conundrum of how the brain achieves its remarkable computational powers on a limited energy budget.

The framework of whole-brain modelling is offering a powerful way to determine causal mechanisms of brain function [7–11]. The best fit to empirical data is offered by whole-brain models using classical coupled Stuart-Landau oscillators to accurately model empirical brain activity [12]. Importantly, these models explain the emergence of metastability, expressing the rich repertoire of brain dynamics needed for complex computation [13, 14] and that removing long-range connectivity in the model significantly reduces the fit to empirical brain data [3, 4].

Interestingly, recent advances in physics have shown so-called quantum-like (QL) interferences at the macroscopic level in systems of non-quantum (classical) coupled oscillators [15–17]. Scholes works within the Växjö interpretation of quantum mechanics [18] to show that QL states are vector-like models for measurement sequences. Furthermore, the eigenstates of classical systems can be combined through so-called ‘QL-bits’ that are the interactions, such as coupled oscillators, between the systems and represent the tensor product of vector spaces, i.e. Hilbert spaces.

Based on this new body of work, we here investigate in the classical macroscopic system of the brain, if non-quantum coupled oscillators can give rise to long-range interactions through QL interference effects. This would be similar, but not equal, to how interference is at the heart of quantum systems characterised by non-locality achieved by long-range interference effects. In particular, we investigate whether QL effects could have an important role to play for the remarkable efficiency of brain computation [19, 20]. We investigate the emergent states produced by cluster-synchronisation of systems of coupled oscillators used in the whole-brain models of empirical brain data [7, 12, 21, 22].

Here results show that QL states are found in empirical human brain activity neuroimaging data. Specifically, we find QL interference effects are amplifying metastability, which can be defined as the switching between different cluster synchronisation states. Metastability has been shown to be a very important feature for brain computation, intuitively expressing the rich repertoire needed for complex computation [13, 14]. The results demonstrate that QL states on the anatomical structural connectivity form a powerful backbone for the efficiency of computation needed for advanced human cognition.

### Quantum-like effects in coupled oscillators

In order to study QL states in the brain, we first study the local level using a network of coupled oscillators with the topology of *k*-regular graphs. We concentrate of the class of directed graphs with no loops and no multiple edges, consisting of *n* vertices and *m* edges, connecting pairs of vertices. This can be described by a connectivity *n* x *n* matrix ***C***, with the weighted *m* edges. The spectrum of eigenvalues of ***C*** characterises the emergent states. Scholes has shown that *k*-regular random graphs, which belong to a class of specific expander graphs, have strong resilience to decoherence when they exhibit a spectrum where the highest (or multiple) eigenvalues are separated from the rest of the spectrum [23]. In *k*-regular random graphs every vertex is connected to *k* edges, i.e. the degree of every vertex is *k*. Let the eigenvalues of this *k*-regular graph be *λ*_0_ ≥ *λ*_1_ ≥ ⋯ ≥ *λ*_*n*−1_. For a *k*-regular graph, the largest eigenvalue is *λ*_0_ = *k* with corresponding eigenvector 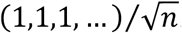. The reason why *k*-regular random graphs are candidates for the emergence of QL states is grounded in the findings from Scholes, using the Alon– Boppana Theorem [24, 25], namely that the spectral gap, *λ*_0_ − *λ*_1_, i.e., the difference between the largest eigenvalue and the second largest eigenvalue, is well-separated from those of the remaining ‘random’ states.

When pruning of the edges in a *k*-regular graph with the probability, *p*_*local*_, the large spectral gap associated with a QL-state is still present for low values of *p*_*local*_. High levels of pruning leading to sufficiently large values of *p*_*local*_, make the spectral gap and consequently the QL state disappear. This is shown in the right panel of **Figure 1A** for a resilient *k*-regular graph with *n* =40, *m* =20 and *p*_*local*_ = 0.2 with its connectivity matrix with the emerging large spectral gap and QL state. In contrast, when constructing a *k*-regular graph using *p*_*local*_ = 0.8 (left panel), the spectral gap disappears. This means that the quantum-likeness of the *k*-regular graph can be regulated with increased levels of pruning.

**Figure 1.**
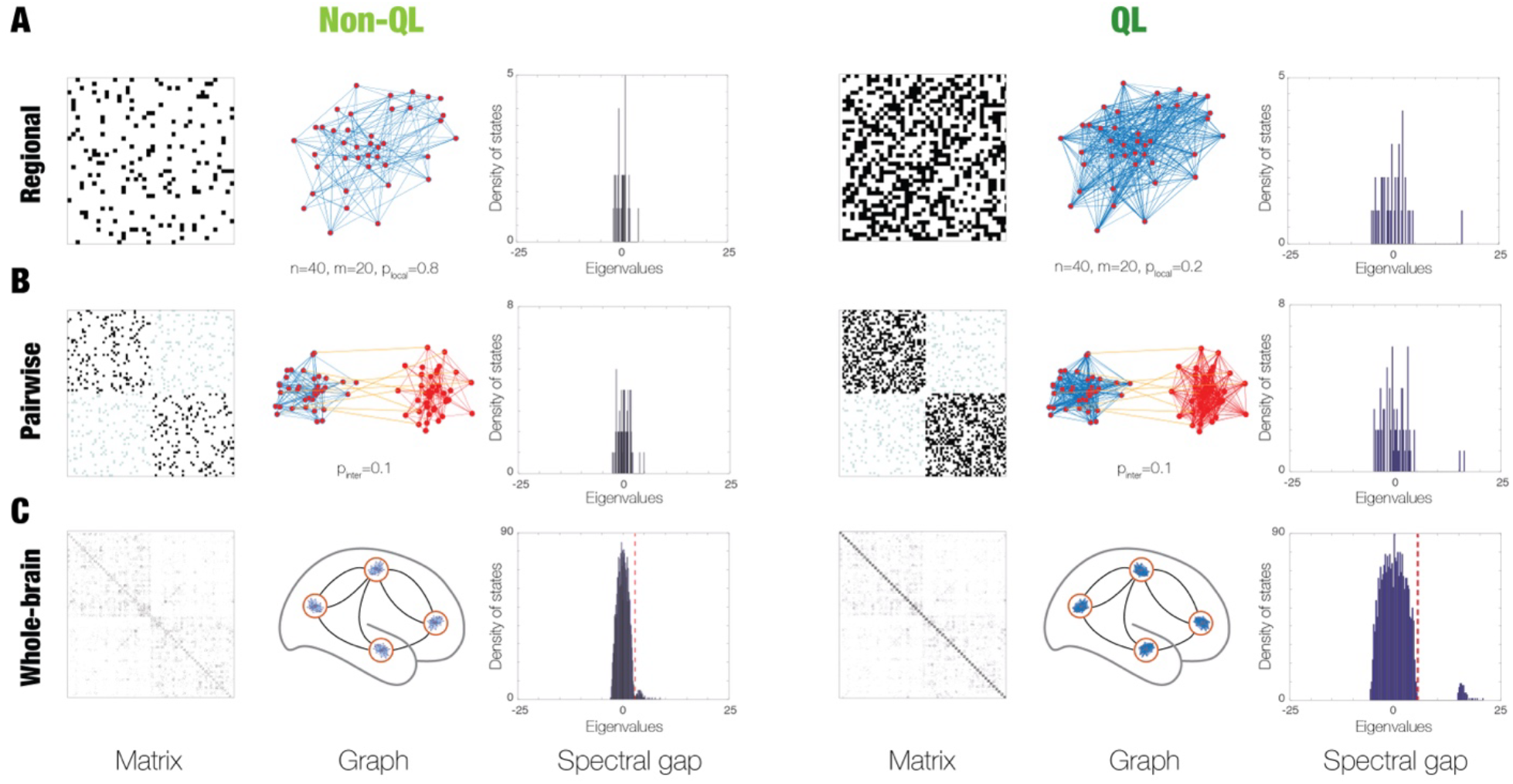
Schematic of the whole-brain modelling with and without quantum-like effects. By using different levels of pruning, coupled oscillators in a k-regular graph can be made to show both non-QL and QL effects. Here we show the construction of whole-brain models with non-QL (left column) and QL (right column) effects at three levels: regional, pairwise and whole-brain, where for each level we show the connectivity matrix, corresponding graph and resulting spectral gap. **A)** For each region (left column) we have a k-regular graph with high levels of pruning, which does not show QL effects, since it does not show a spectral gap in the eigenvalues of the matrix. In contrast (in the right column) for low levels of pruning, this leads to a spectral gap. **B)** A QL bit is typically defined pairwise as the interconnection of two k-regular graphs. Left column show the results of using a coupling of non-QL graphs with the corresponding matrix and spectral gap. Comparing this with the results of using QL graphs (in the right column), this gap is significantly larger. **C)** Defining a network of networks with distributed bits across the whole brain leads to a significantly larger spectral gap when using QL regional graphs as the building blocks for each brain region.

### Hilbert space

Similar to genuine quantum states, QL states can be expressed as vectors in a Hilbert space and any interference effect can thus be described in the same way. Briefly, a Hilbert space ℋ is a complete vector space over a field ℤ. Here we use a real field with the inner product ⟨*x*, *y*⟩, thereby generalising the Cartesian dot product. So, for any vectors *x*, *y*, *z* in ℋ and with scalars *α* ∈ ℤ, the following equalities are satisfied:

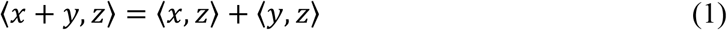

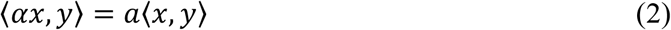

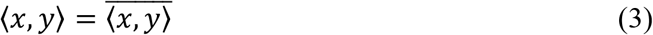

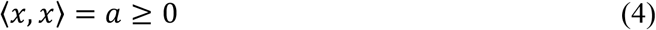

where overbar signifies the complex conjugate. The norm is defined in terms of the inner product: 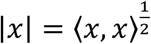. Scholes showed that QL states emerging from the underlying graphs satisfy the following axioms [23]: A) Graphs will show an emergent state distinguishable in the eigenvalue spectrum. B) The graphs are robust and stable to disorder. C) The QL state is a two-state system. In a network of oscillators sustained by the underlying graphs, the two states correspond to synchronisation or no synchronisation. Similar to a real quantum state, interactions between different QL networks of oscillators will be able to produce interference superposition.

### QL bit

Khrennikov and colleagues [18] have shown how classical systems can produce probabilistic outcomes similar to the ones found in quantum mechanical systems. Following this line of research, Scholes proposed that a network of coupled oscillators can produce QL states, which are robust to decoherence and show interference superposition effects as expressed in a Hilbert space. In other words, a network of coupled oscillators is a perfect implementation of QL states. Combining two QL networks of coupled oscillators produces a QL bit [15]. Through cluster synchronisation, such QL bits could generate two separate QL states that can interfere and with superposition ruled by a Hilbert space. **Figure 1B** (right panel) shows a QL bit consisting of two coupled *k*-regular graphs (*n* = 40, *m* = 20 and *p*_*local*_ = 0.2) whose vertices are interconnected with a probability of *p*_*inter*_ = 0.1. As before, the local weights are 1, while the inter-network weights are 0.17. The two QL states emerge due to the large underlying spectral gap, creating the possibility of building a large classical system with quantum superposition implemented through cluster synchronisation. In contrast, by manipulating the *k*-regular graphs through increasing the pruning probability *p*_*local*_ = 0.8, these states do not show QL states (shown in **Figure 1B**, left panel).

### Whole-brain models

having established that *k*-regular networks with coupled oscillators show interference superposition, we hypothesise that implementing these in a whole-brain model of empirical human brain activity will show that the brain computation is quantum-like. The dynamics of each brain region will be governed by a *k*-regular network with coupled oscillators connected with the anatomical structural connectivity and as a result this distributes QL bits across paired brain regions and thus distributed, connected and overlapping across the whole brain (see middle panels of **Figure 1C** for non-QL and QL cases). Importantly, we used coupled Stuart-Landau (SL) oscillators (i.e., as the normal form of a supercritical Hopf bifurcation) since we have shown that these provide excellent fitting of whole-brain activity [7, 12, 21] given that they are biophysical realistic and equivalent to an exact dynamic mean field model of quadratic integrate-and-fire neurons [26]. Furthermore, and crucially, the best working point of SL oscillators are at the edge of the bifurcation, which permitted us to create a linearisation of the model [27]. This makes possible for the first time to create the necessary large-scale analytical models of networks of networks to describe the empirical human neuroimaging data.

Briefly, for a local brain region, we use a network of coupled oscillators connected by *k*-regular network with *n* = 40 and *m* = 20, with two different levels of pruning, where QL effects emerge for low levels of pruning (*p*_*local*_ = 0.2), and disappear for high levels of pruning (*p*_*local*_ = 0.8). All brain regions are coupled using the anatomical structural connectivity matrix ***C***, expressing the fibre density obtained with diffusion magnetic imaging (MRI) and normalised to 1. The SL oscillators between paired brain regions (*i*, *j*) are connected with the probability of *p*_*inter*_ = 0.1 and scaled by *C*_*ij*_. Consequently, this build a large-scale network of networks 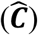, that consists of the full brain network of *N* regions and where each region has a *k*-regular network with *n* regions, leading to *M* = *N* ∗ *n*. **Figure 1C** shows two examples of these large matrices 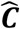 (with dimensions of *M* × *M*) with high pruning on the left (non-QL) and low pruning on the right (QL). **Figure 2** schematises the elements used for the whole-brain model. Mathematically, the whole-brain dynamics are defined by

**Figure 2.**
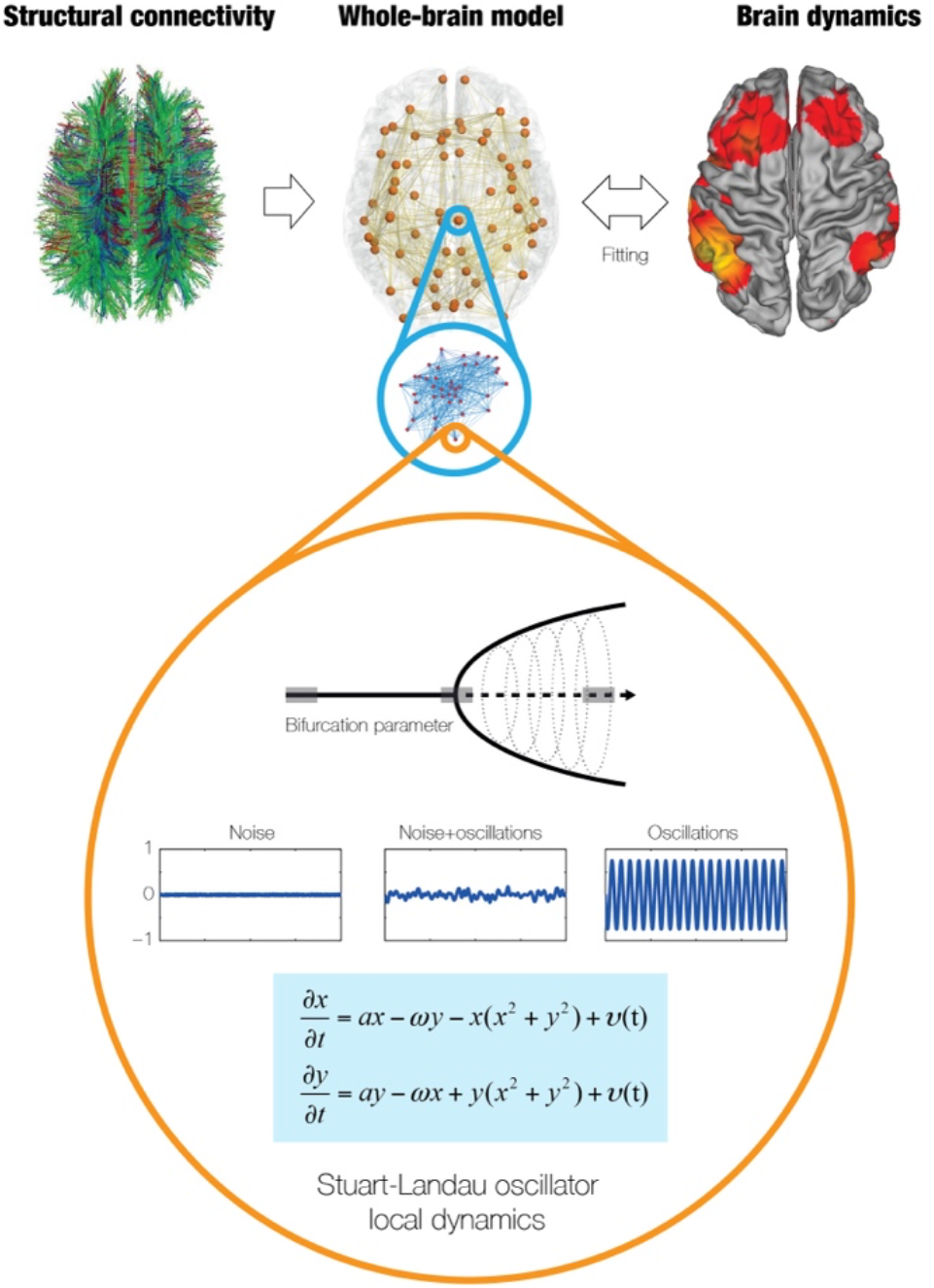
Whole-brain model with k-regular graphs in each brain region. The dynamics of each brain region is governed by a k-regular network with coupled Stuart-Landau (SL) oscillators connected with the anatomical structural connectivity and as a result this distributes QL bits across paired brain regions and thus distributed, connected and overlapping across the whole brain.

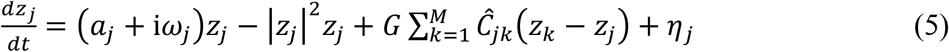

where for the oscillator in region *j*, the complex variable *z*_*j*_ denotes the state (*z*_*j*_ = *x*_*j*_ + i*y*_*j*_), *η*_*j*_ is additive uncorrelated Gaussian noise with variance *σ*^2^ (for all *j*), *ω*_*j*_ is the intrinsic node frequency, and *a*_*j*_ is the node’s bifurcation parameter. Within this model, the intrinsic frequency *ω*_*j*_ of each node is in the 0.008–0.08Hz band. The intrinsic frequencies were estimated from the empirical data, as given by the averaged peak frequency of the narrowband blood-oxygen-level-dependent (BOLD) signals of each brain region. For *a*_*j*_ > 0, the local dynamics settle into a stable limit cycle, producing self-sustained oscillations with frequency *ω*_*j*_/(2*π*). For *a*_*j*_ < 0, the local dynamics present a stable spiral point, producing damped or noisy oscillations in the absence or presence of noise, respectively. The fMRI signals were modelled by the real part of the state variables, i.e., *x*_*j*_ = Real(*z*_*j*_). The fitting of the whole-brain model is accomplished by finding the optimal fit of the global coupling parameter (global conductivity), *G*, by minimising the elementwise quadratic error between the functional connectivity matrices (Pearson correlations between the fMRI timeseries in each brain region) of the model (***FC***^*model*^) and empirical (***FC***^*emp*^) data.

This large-scale optimisation is made possible by the linearisation of the SL oscillator [27]. Proximity of the SL oscillator to criticality has been shown to provide the best working point for fitting whole-brain neuroimaging dynamics. This happens at the brink of the bifurcation, i.e. with *a*_*j*_ slightly negative but very near to zero (usually *a*_*j*_ = −0.02) [12]. Briefly, we can estimate the functional correlations of the whole-brain network using a linear noise approximation (LNA). Hence, the dynamical system of *N* nodes (**Equation 5**) can be re-written in vector form as:

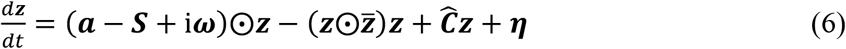

where ***z*** = [*z*_1_, … , *z*_*M*_]^*T*^, ***a*** = [*a*_1_, … , *a*_*M*_]^*T*^, ***ω*** = [*ω*_1_, … , *ω*_*M*_]^*T*^, ***η*** = [*η*_1_, … , *η*_*M*_]^*T*^ and ***S*** = [*S*_1_, … , *S*_*M*_]^*T*^ is a vector containing the strength of each node, i.e. *S*_)_ = ∑_*j*_ *Ĉ*_*ij*_. The superscript [ ]^*T*^represents the transpose, 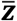 is the complex conjugate of ***z*** and ⨀ is the Hadamard element-wise product. As such, the equation describes the linear fluctuations around the fixed point ***z*** = 0, which is the solution of 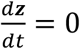. Discarding the higher-order terms 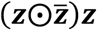 and separating the real and imaginary parts of the state variables, the evolution of the linear fluctuations follows a Langevin stochastic linear equation:

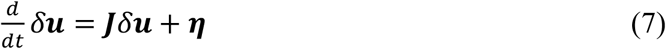

where the 2*M*-dimensional vector *δ****u*** = [*δ****x***, *δ****y***]^***T***^ = [*δx*_1_, … , *δx*_*M*_, *δy*_1_, … , *δy*_*M*_]^*T*^ contains the fluctuations of real and imaginary state variables. The 2*M* × 2*M* matrix ***J*** is the Jacobian of the system evaluated at the fixed point, which can be written as a block matrix

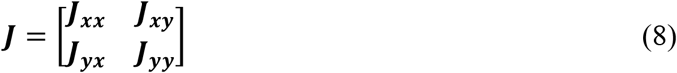

where ***J***_***xx***_, ***J***_***xy***_, ***J***_***yx***_, ***J***_***yy***_ are *M* × *M* matrices given as ***J***_***xx***_ = ***J***_***yy***_ = diag(***a*** − ***S***) + ***C*** and ***J***_***xy***_ = −***J***_***yx***_ = diag(***ω***), where diag(***v***) is the diagonal matrix whose diagonal is the vector ***v***. Please note that this linearisation is only valid if ***z*** = 0 is a stable solution of the system, that is if all eigenvalues of ***J*** have negative real parts. To compute the covariance matrix ***K*** = ⟨*δ****u****δ****u***^***T***^⟩, one can begin by writing **Equation 6** as *dδ****u*** = ***J****δ****u****dt* + *d****W***, where *d****W*** is an 2*M*-dimensional Wiener process with covariance ⟨*d****W****d****W***^***T***^⟩ = ***Q****dt*, where ***Q*** is the noise covariance matrix, which is diagonal if the noise is uncorrelated. Using Itô’s stochastic calculus, we get *d*(*δ****u****δ****u***^***T***^) = *d*(*δ****u***)*δ****u***^***T***^ + *δ****u****d*(*δ****u***^***T***^) + *d*(*δ****u***)*d*(*δ****u***^***T***^). Noting that ⟨*δ****u****d****W***^***T***^⟩ = 0, taking the expectations and keeping terms to first order in the differential *dt*, we obtain:

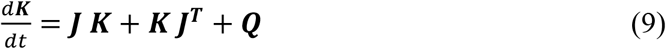

Hence, the stationary covariances (for which 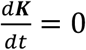) can be obtained by solving the following analytic equation:

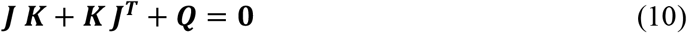

**Equation 10** is a Lyapunov equation that can be solved using the eigen-decomposition of the Jacobian matrix [28]. We obtained the simulated functional connectivity matrix ***FC***^*model*^ by reducing (through averaging across the *n* in each brain region) the functional connectivity obtained from the first *M* rows and columns of the covariance ***K***, which corresponds to the real part of the dynamics, and thus the BOLD fMRI signal.

### QL effects in empirical brain dynamics

Figure 3. shows the fitting of whole-brain models with non-QL and QL local dynamics (with and without distributed spectral gaps as seen in **Figure 1C**, compare the spectral gaps shown in right panels for non-QL and QL). As can be seen, there is a very significant difference in the level of fitting of large-scale human neuroimaging resting state data from over 1000 healthy participants (see Appendix) in terms of correlation (**Figure 3A**) and quadratic error (**Figure 3B**) between the simulated and empirical FC matrices. **Figure 3C** demonstrates that the differences in fitting are driven by significant difference in the underlying spectral gaps shown in the violin plots. The values are calculated by summing all the spectral gap differences by summing between eigenvalues larger than the red line (threshold) in the histograms of the spectral gaps for non-QL and QL shown in **Figure 1C**. We calculated (*λ*_0_ − *λ*_1_) + (*λ*_1_ − *λ*_2_) + ⋯) over this threshold for both non-QL and QL. **Figure 3D** shows renderings of the regional spectral gaps on the surface of the human brain for both non-QL and QL. Each region contains the sum of the pairwise spectral gap between this region and all other regions. This shows the clear differences between non-QL and QL. Importantly, for the QL case, there is large heterogeneity in the spectral gaps across brain regions suggestive of differences in quantum-like effects across the brain.

### Energy cost of QL computation

**Figure 3E** quantifies the energy consumption linked with the whole-brain modelling based on the recent COCO framework [20], which combines stochastic thermodynamics with whole-brain modelling for calculating analytically energy consumption, entropy production and information processing from neuroimaging data (see Appendix). Excitingly, the results show a very significant drop in energy consumption for QL compared to non-QL. The quantum-like nature of the human brain demonstrated here for the first time could a driving reason for why the human brain is so energy efficient and uses much less energy than other species [20].

**Figure 3.**
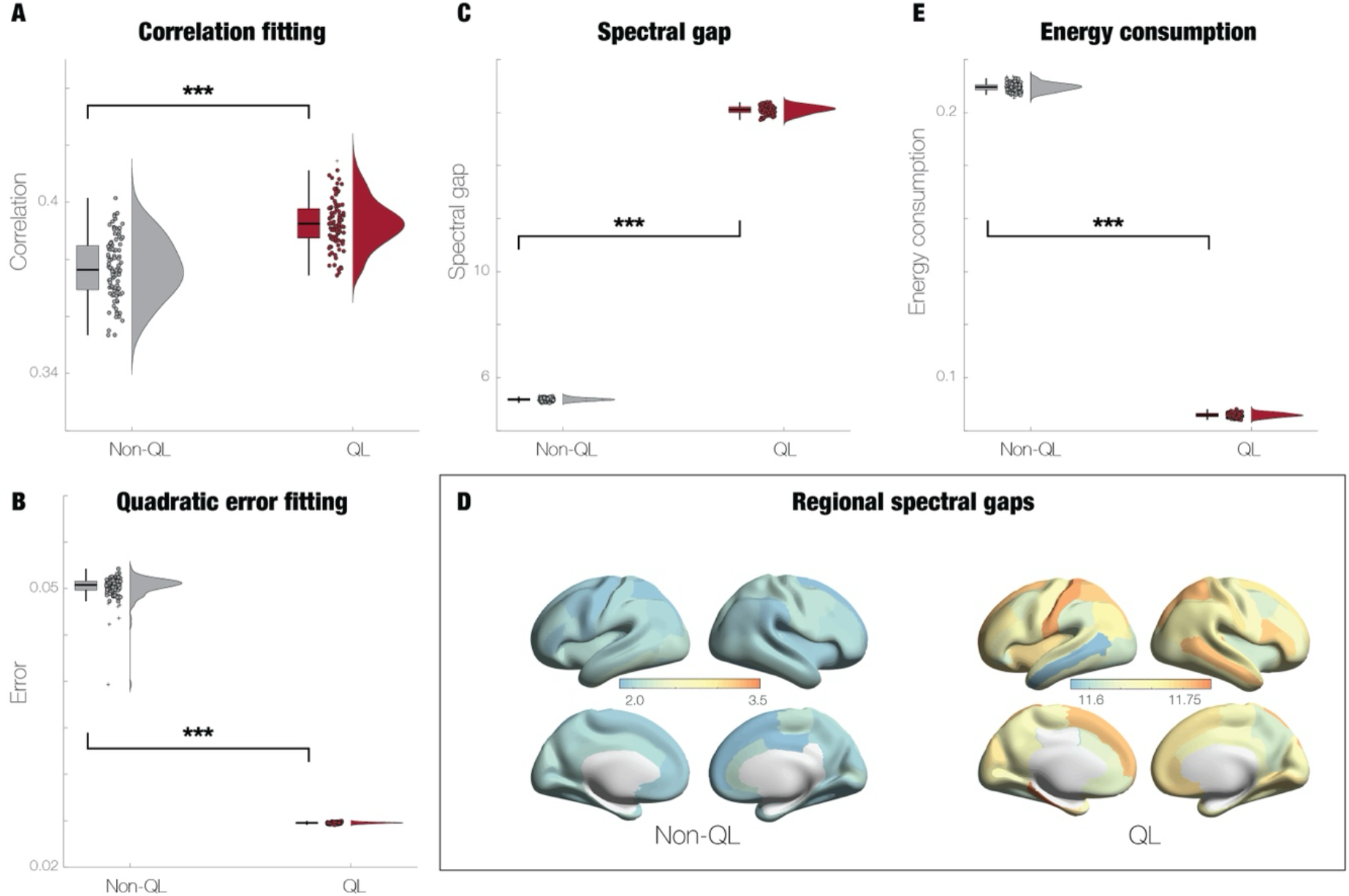
Evidence of quantum-like (QL) computation in the human brain. **A)** Fitting of whole-brain models with non-QL and QL local dynamics show significant differences in the level of fitting of large-scale human neuroimaging resting state data in terms of correlation the simulated and empirical FC matrices. **B)** The quadratic error fitting is also significantly better for the QL whole-brain model. **C)** Significant differences in the underlying spectral gaps shown in the violin plots. **D)** Renderings of the regional spectral gaps on the surface of the human brain for both non-QL and QL.

### Special anatomical brain structure

previous research has pointed to a very significant role of a unique feature of brain anatomy [3, 4], namely the rare long-range exceptions to the exponential distance rule [5, 6]. Kennedy and colleagues used retrograde tract tracing in non-human primates to describe this rule which explains the local connectivity of the brain solely in terms of the geodesic distance between points on the cortical surface. Given that the anatomy of the human brain exhibits QL effects, we investigated if removing the rare long-range exceptions would significantly affect the QL effects. In order to do so, we constructed individual whole-brain QL regime models fitting the empirical data from each of the 1003 human participants. We did this for whole-brain models with and without rare long-range exceptions over 80 mm. In order to individualise the models we used the principle of creating the generative effective connectivity for each individual [29] (see Appendix). We used the spectral gap as a signature of QL effects and **Figure 4A** shows the significant decrease in spectral gaps when the long-range exceptions are removed. This clearly demonstrates that the long-range exceptions amplify the quantum-like effects of human computation. Equally, this allowed us to establish a correlation between metastability (richness of repertoire, see Appendix) and the QL effects (measured by spectral gap). **Figure 4B** shows the correlations between metastability and spectral gap between the individualised QL whole-brain models with and without long-range exceptions. As can be seen, the QL whole-brain models with long-range exceptions have a stronger correlation than without. This further demonstrates that metastability (as a measure of cluster synchronisation) underlies the QL spectral gap and that this is amplified by the unique long-range exceptions to the exponential distance rule.

**Figure 4.**
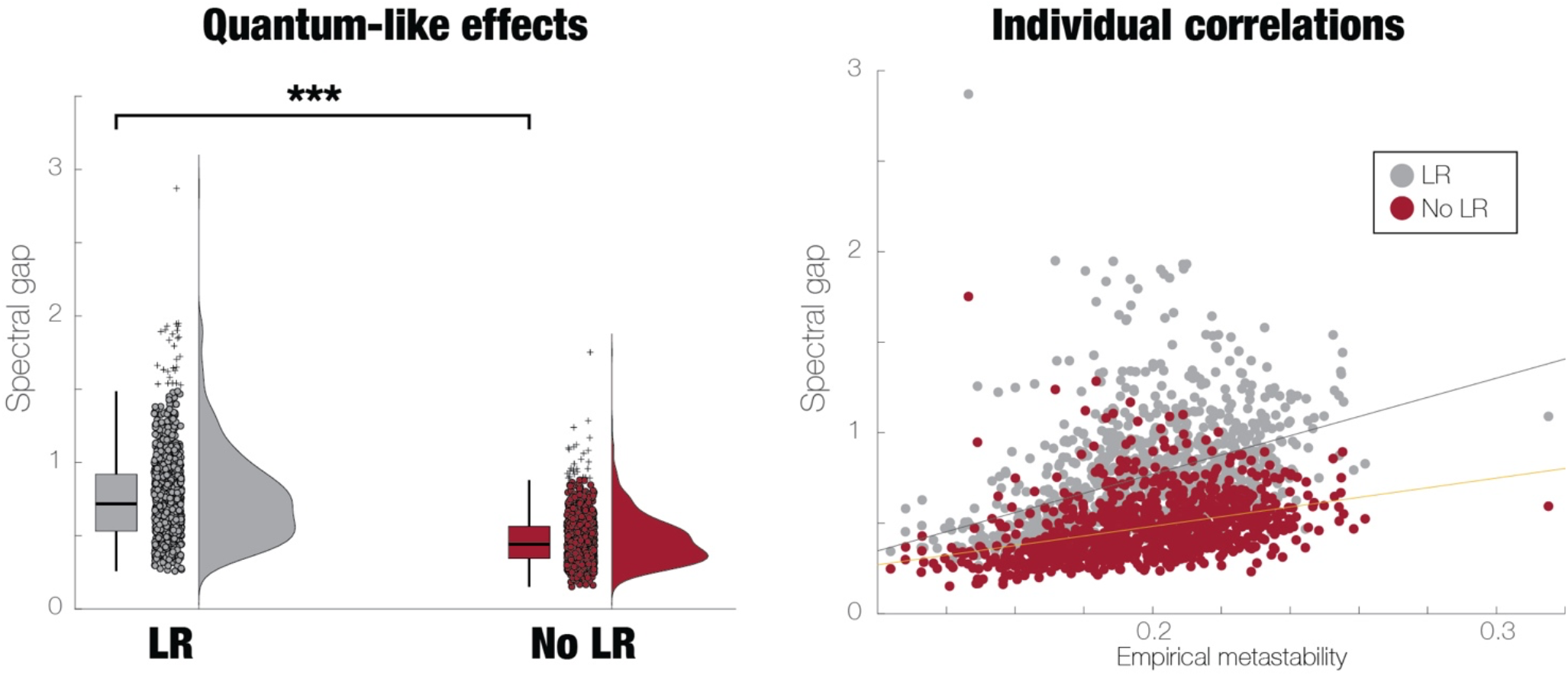
Evidence for the importance of long-range connections amplifying quantum-like (QL) computation in the human brain. **A)** There is a significant decrease in spectral gaps when the long-range exceptions (>80mm) are removed from the underlying structural anatomical connectivity, showing that long-range exceptions amplify the quantum-like effects of human computation. **B)** As seen from the scatter plot, there is a significantly stronger correlation between metastability and the QL effects (measured by spectral gap) between the individualised QL whole-brain models with and without long-range exceptions.

## Discussion

This is the first evidence that the human brain exhibit quantum-like (QL) dynamics, demonstrated using whole-brain modelling with or without QL effects. Specifically, we show that whole-brain modelling using non-quantum (classical) coupled oscillators fits human brain activity significantly better when using a manipulation with using either QL versus non-QL local dynamics in brain regions, where the QL effects can be seen from the spectral gaps. Furthermore, we show that QL whole-brain models are significantly more energy efficient than non-QL models. Importantly, the QL effects are amplified by the special brain architecture with long-range exceptions to the predominant local exponential distance rule [5, 6]. These results strongly suggest that the QL dynamical effects demonstrated in human brain activity could in fact be the computational backbone of human cognition.

Supporting this conjecture is the fact that the long-range connectivity is strongest in humans compared to other species. When long-range exceptions to the structural connectivity are removed from the whole-brain model, there is a significant decrease in the QL effects as indexed by the spectral gaps. This demonstrates that long-range connectivity provides an amplification of quantum-like dynamics of human brain activity. As such, this provides more evidence that the rare long-range exceptions to the anatomical exponential distance rule of human brain connectivity are crucial for human cognition [3, 4].

Further bolstering this link to the underlying human brain architecture, but now demonstrating the importance for functional dynamics, we demonstrated a correlation between QL effects and metastability (richness of repertoire). QL whole-brain models with long-range exceptions have a stronger correlation than whole-brain models with non-QL dynamics. This suggests that metastability, which is a measure of variability of cluster synchronisation, is expressing dynamically the underlying QL spectral gap and is amplified by the unique long-range connectivity. More generally, it is important given the crucial role of metastability as a very useful concept from dynamical systems, which can be used to capture the balance between integration and segregation needed for healthy brain dynamics [14, 30, 31] and has been shown to be a sensitive marker of disease [32, 33].

In fact, the efficiency in computation achieved by QL in the human brain is linked to the increase in the richness of repertoire as captured by metastability (rich switching in different cluster synchronisation patterns). Here the results suggest novel insights into how the richness of repertoire in brain dynamics is supported by QL effects, promoting the switching between different patterns of cluster synchronisation. These results may also be related to the relevance the field of quantum cognition [34, 35] that explains paradoxes in human cognition using an alternative probabilistic framework drawn from the mathematical structure of quantum theory.

Equally, the findings presented here are likely related to the effects from quantum mechanics which has revolutionised the way we understand the microscopic world, especially for describing non-local entanglement as the core of quantum states [36–38]. It has been shown that the probability laws governing these states and in particular the interference effects are crucial for effective computation found in quantum information science [39–42] but quantum decoherence makes it difficult to construct robust quantum computing [43]. This link could be relevant given that QL effects are found in the brain which is a classical macroscopic system. Importantly, the QL effects are robust and do not exhibit the fragility of the well-known process of quantum decoherence [43]. As a result, in future the robust QL effects demonstrated in the brain could perhaps be important for the emergence of true macroscopic quantum computing.

Overall, the findings reported here are important since they have shed new light on how the human brain is able to sustain computation on the remarkably small energy budget of the human brain continuously running on around 20 watts of power [1, 2]. This is in stark contrast to current artificial intelligence, where a typical large high-performance computing cluster use up to six orders of magnitude more power to operate at typically 2 megawatts. Given that QL whole-brain models fit human brain activity better than non-QL models, the energy efficiency of QL models could be a very important finding. This could offer a potential explanation of why the cost of brain cognition is so much more efficient in biological brains than in current computing systems.

